# Computational mechanisms underlying motivation to earn symbolic reinforcers

**DOI:** 10.1101/2023.10.11.561900

**Authors:** Diana C. Burk, Craig Taswell, Hua Tang, Bruno B. Averbeck

## Abstract

Reinforcement learning (RL) is a theoretical framework that describes how agents learn to select options that maximize rewards and minimize punishments over time. We often make choices, however, to obtain symbolic reinforcers (e.g. money, points) that can later be exchanged for primary reinforcers (e.g. food, drink). Although symbolic reinforcers are motivating, little is understood about the neural or computational mechanisms underlying the motivation to earn them. In the present study, we examined how monkeys learn to make choices that maximize fluid rewards through reinforcement with tokens. The question addressed here is how the value of a state, which is a function of multiple task features (e.g. current number of accumulated tokens, choice options, task epoch, trials since last delivery of primary reinforcer, etc.), drives value and affects motivation. We constructed a Markov decision process model that computes the value of task states given task features to capture the motivational state of the animal. Fixation times, choice reaction times, and abort frequency were all significantly related to values of task states during the tokens task (n=5 monkeys). Furthermore, the model makes predictions for how neural responses could change on a moment-by-moment basis relative to changes in state value. Together, this task and model allow us to capture learning and behavior related to symbolic reinforcement.

**Significance statement:** Symbolic reinforcers, like money and points, play a critical role in our lives. Like rewards, symbolic reinforcers can be motivating and can even lead to compulsive behaviors like gambling addiction. However, we lack an understanding of how symbolic reinforcement can drive fluctuations in motivation. Here we investigated the effect of symbolic reinforcers on behaviors related to motivation during a token reinforcement learning task, using a novel reinforcement learning model and data from five monkeys. Our findings suggest that the value of a task state can affect willingness to initiate a trial, speed to choose, and persistence to complete a trial. Our model makes testable predictions for within trial fluctuations of neural activity related to values of task states.

## Introduction

In most decision-making contexts, the objective is to maximize rewards and minimize punishments over time. In some situations, rewards are symbolic, such as money or points, in which case they can be exchanged for primary rewards, such as food or drink, in the future. Past studies have shown that animals and humans will work for symbolic reinforcers, and symbolic reinforcers can drive learning and therefore motivate behavior (Hackenberg, 2009, 2018).

Motivation is a process that invigorates behavior in the present to reach rewards in the future (Berridge, 2004; Berke, 2018; O’Reilly, 2020). Motivation can be studied in the context of reinforcement learning (RL). Learning builds predictions of choice outcomes that can be used to direct future behavior (Sutton and Barto, 1998). N-armed bandit tasks are often used to study RL in animals. These tasks can be modeled with Rescorla-Wagner (RW) RL models (Recorla, 1972), because the choices lead probabilistically, but immediately, to a primary reinforcer (Bartolo and Averbeck, 2020; Beron et al., 2022). However, one can also use symbolic reinforcers, for example, tokens or money to drive learning (Jackson, 1996; Kirsch et al., 2003; Seo and Lee, 2009; Delgado, Jou and Phelps, 2011; Taswell et al., 2018; Taswell et al., 2021; Falligant and Kranak, 2022; Yang, Li and Stuphorn, 2022; Taswell et al., 2023). In these tasks, subjects learn to make choices to obtain tokens, which can be exchanged in the future for primary reinforcers. Token based learning tasks set up a distinction between two types of cues that predict rewards in different ways. Specifically, tokens predict rewards on long-time scales, deterministically, whereas choice cues predict tokens. Furthermore, the relation between cues and tokens must be learned. Such tasks involving symbolic reinforcement cannot be accurately captured with current RL models, because the distinction between rewards and symbolic reinforcers cannot be made explicit.

Thus, we developed a Markov Decision Process (MDP) model to characterize the value of symbolic reinforcers, and the computational mechanism that links cues through tokens to rewards. The MDP also allows us to model multiple factors that drive value, including the time to reach primary rewards, and the probability of obtaining additional rewards in the future. We can therefore use the model to establish which factors most strongly drive behavior. To establish the validity of the model, beyond predicting learning which can be done with RW-RL models, we examined the relationship between the state value (i.e. the expected discounted sum of future rewards) and behavioral measures associated with motivation. Five monkeys performed a task where they learned to select images that led to gaining or losing tokens. The tokens were later exchanged for juice rewards. To examine the ability of the model to capture motivation, we conducted regressions between state value and change in state value and three behaviors linked to motivation. To demonstrate the effect of each task dimension included in the model, we performed marginalization analyses, where each feature was removed and the analyses were repeated. These analyses demonstrated that all features that drove value in the model contributed to motivation. Taken together, our results make predictions for how neural activity might evolve in reinforcement learning circuits during a task involving symbolic reinforcement.

## Materials and Methods

### Subjects

The subjects included three male and two female rhesus macaques with weights ranging from 6 to 11 kg. Four monkeys were used as control monkeys in a previous study (Taswell et al., 2018). One additional monkey was a naïve monkey whose first task was the tokens task. For the duration of collecting behavioral data, monkeys were placed on water control. On testing days, monkeys earned their fluid from performance on the task. Experimental procedures for all aspects of the study were performed in accordance with the Guide for the Care and Use of Laboratory Animals and were approved by the National Institute of Mental Health Animal Care and Use Committee.

### Experimental Design

We conducted post hoc analyses on previously published data (Taswell et al., 2018) and data from one additional subject. We use data from one variant of the tokens task, previously called Stochastic Tokens with Loss (referred to as TkS).

The images used in the task were normalized for luminance and spatial frequency using the SHINE toolbox for MATLAB (Willenbockel et al., 2010). Image presentation was controlled by PC computers running Monkeylogic toolbox (Version 1.1) for MATLAB (Asaad and Eskandar, 2008; Hwang, Mitz and Murray, 2019). Eye movements were monitored using the Arrington ViewPoint eye-tracking system (Arrington Research).

### Stochastic Tokens Task with Gains and Losses

Blocks consisted of 108 trials that used four novel images that had not been previously presented to the animal. Each image was associated with a token outcome (+2, +1, −1, −2), such that if that image was chosen, the animal gained or lost the corresponding number of tokens 75% of the time and received no change in tokens 25% of the time (**Fig. 1A**).

**Figure 1.**
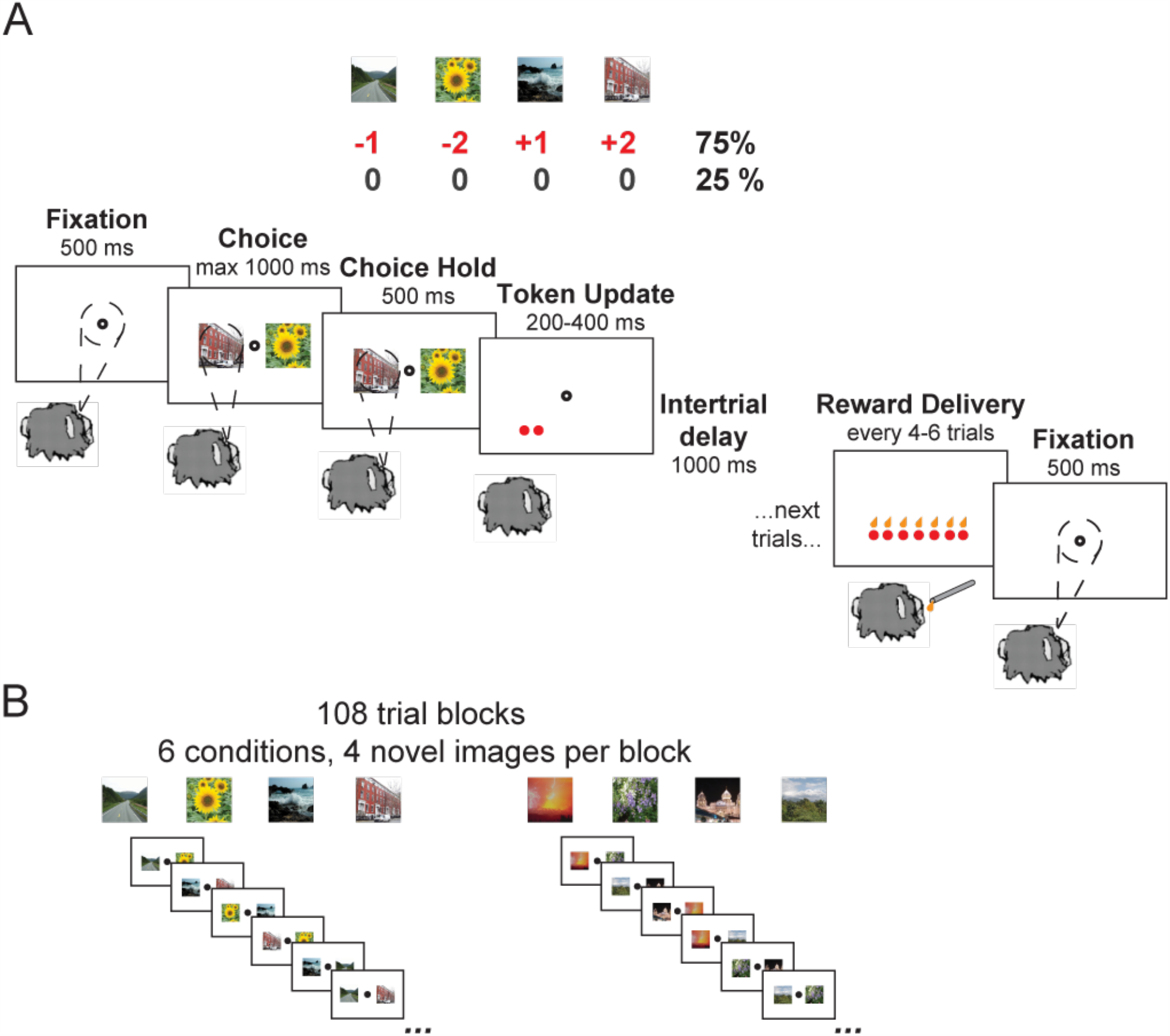
Overview of the Tokens task with stochastic rewards. **(A)** Flow of single trial. First the monkey fixates on the screen and is required to hold fixation for 500 ms. Two cue images appear on either side of fixation, each of which is associated with gaining or losing tokens (+2, +1, -2, -1). The monkey must make a saccade to one of the images and hold their gaze for 500 ms. After a successful hold, the number of tokens associated with the chosen option appears on the screen 75% of the time, and 25% of the time, nothing changes. After a 1000 ms intertrial delay, the next trial begins. Every four to six trials, tokens were exchanged for juice drops 1:1 and the monkey started the subsequent trial with zero tokens. **(B)** The monkeys learned through trial and error which visual images were associated with gaining tokens and which images were associated with losing tokens. Four new images were presented every block of 108 completed trials. Each pair of cues (six total) was seen 18 times (nine Left/Right, nine Right/Left). Image credit: Wikimedia Commons (scene images).

On each trial, monkeys had 2000 ms to acquire a fixation spot at the center of the screen and were required to hold fixation for 500 ms. After monkeys held central fixation, two of the four possible images would appear to the left and the right of the fixation point. The animal had 1000 ms to choose one of the images by making a saccade to an image and hold their gaze on the image for 500 ms to indicate their choice. If the monkey moved his eyes outside the fixation window during fixation, did not choose a cue, or did not hold the cue long enough, the trial was aborted and repeated immediately. After a successful hold of gaze on a choice, tokens associated with the image were then added or subtracted from their total count, represented by circles at the bottom of the screen. Note that the animals could not have fewer than zero tokens. After an intertrial interval of 1000 ms, the next trial would begin with the accumulated tokens visible on the screen the entire time. Every four to six trials, tokens were exchanged 1:1 for juice drops. During this cashout epoch, one drop of juice was delivered and a token disappeared, until all tokens were gone. The animal did not choose when to cash out, rather the probability of exchanging tokens for juice drops was a uniform distribution over four to six trials.

There were six cue conditions in the task, defined by the possible pairs of the four images. The conditions within a block were presented pseudorandomly, such that the animals saw each condition twice (same images, opposite sides) every 12 trials before seeing any condition a third time. This prevented strings of trials with loss v. loss that could lead to abberant behaviors. At the end of each 108-trial block, we introduced four new images and the animals restarted learning associations between the pictures and the token outcomes (**Fig. 1B**). The animals completed approximately 9 blocks of images per session, and approximately 20 sessions from each animal were used for subsequent analyses.

### Model Framework

We modeled the Tokens task using a Markov decision process with partially observable states (POMDP). To fit the POMDP to each animal’s behavior, we leveraged the Rescorla Wagner Reinforcement Learning Model (RW-RL) to calculate the average values of each of the four cue images and to verify the validity of our MDP results for fitting choice probability curves across the cue conditions. Details of this process are in the following sections.

### Rescorla Wagner Reinforcement Learning (RW-RL) Model

We used a variant of the RW-RL model as was previously used to model the tokens task (Taswell et al., 2018).

We fit a Rescorla-Wagner value update equation given by the following:

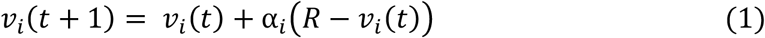

where the variable *v*_*i*_ is the value estimate for cue option *i, R* is the change in the number of tokens that followed the choice in trial *t*, and α_*i*_ is the cue-dependent learning rate parameter. In past work on this data, the model with a separate learning rate parameter for each cue was found to be the best RW-RL model fit to the data. Thus, we continued using this formulation of the RW-RL for these analyses, although the results described in this study are not contingent on this choice.

The value computed in Eq. 1 were then used to compute choice probabilities for each cue pair using the softmax function:

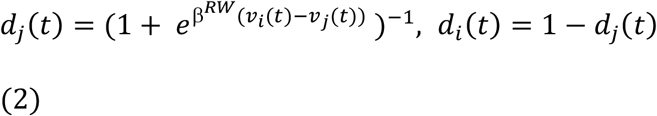

where β, is the consistency choice parameter, fit across all six cue conditions, and *i* and *j* are the two choice options. We then maximized the likelihood of the animal’s choices, *D*, given the parameters, using the cost function:

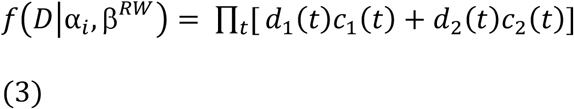

where *d*_1_(*t*) is the choice probability value for option 1 on trial t and *c*_1_(*t*) and *c*_2_(*t*) are indicator variables that take on a value of 1 if the corresponding option was chosen and 0 otherwise. This model was fit across blocks in each session for each monkey to give one set of fit parameters for each session.

Mean cue values as a function of learning trial in each block from the RW-RL model were used to generate transition probabilities for the MDP discussed in detail below. To extract mean cue values, all *v*_*i*_ for a single cue were averaged across sessions. This produced four curves that reflected the change in cue value across trials for each animal (**Fig 2A, 2B**).

**Figure 2.**
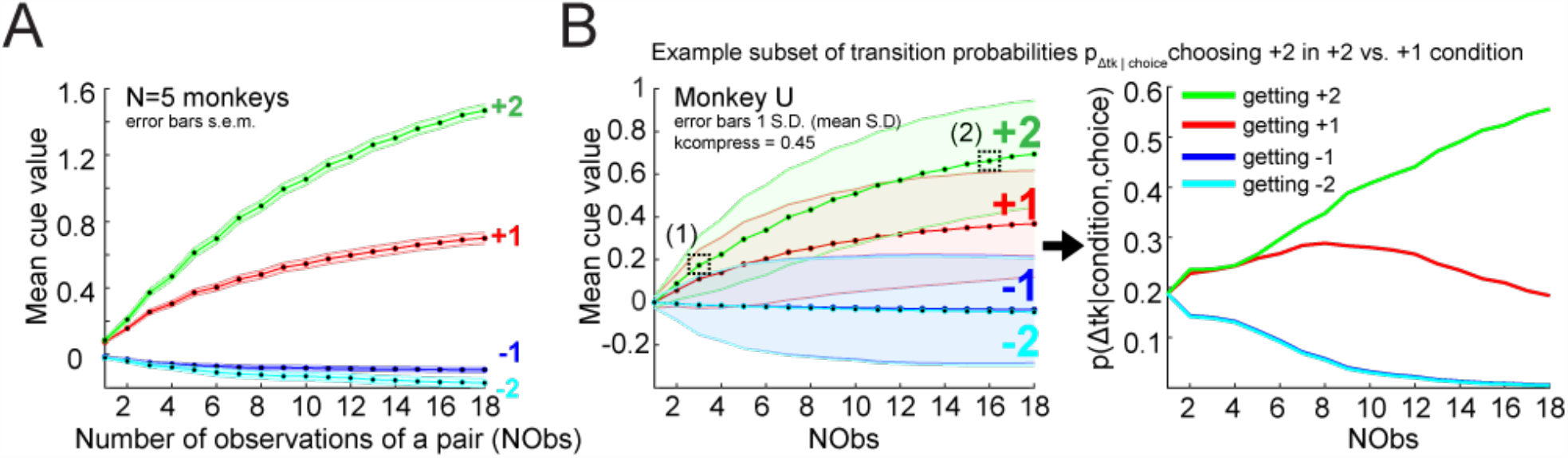
Mean cue values extracted from the RW-RL model and transition probabilities for change in tokens given a cue image selection. **(A)** Mean cue values across number of observations of a cue pair (NObs) are plotted for each of the four cues in the stochastic tokens task (error bars s.e.m. across monkeys). The +2 curve is the highest, followed by the +1 curve, reflecting how the animals learned to select the +2 and +1 image cues. -2 and -1 curves are similar, reflecting that the animals did not learn the value differences between the -2 and -1 cue options. The mean values were extracted from the RW-RL model fits to each monkey. Each data point is an average of the three conditions in which the cue was observed. For example, for the +2 curve, a single data point would be the average value of the +2 cue value from the conditions +2 v +1, +2 v -2, +2 v -1. **(B)** Example set of curves from Monkey U with scaling applied to the variance (see Methods) and scaling parameter fit during the MDP fitting process and derivation of a subset of transition probabilities. Left: (1) +2 cue value highlighted early in learning at trial 3. (2) +2 cue highlighted late in learning at trial 16. Error bars are the mean variance across blocks and solid lines show the cue values with the parameterized scaling factor. Right: Plot of a subset of transition probabilities derived from mean value curves for the +2 v +1 condition and choice of +2 for each possible token outcome. p(Δtk=0|choice +2 cue) is always 0.25 and is not shown.

### Markov Decision Process (MDP) Model

The MDP model computes the value of each task state. Task states were defined by four features of the task: number of tokens (NTk), trials since cashout (TSCO), task epoch (TE), and number of observations of a cue pair (NObs). The state space included all possible combinations of these features across a single block of trials, such that the bounds for each feature were: NTk: 0-12, TSCO: 1-6, TE: 1-10 (which included fixation, 6 cue conditions, token outcome, cashout, intertrial interval), NObs: 1-18. Using NObs as a feature allowed us to avoid having to track each time a cue was shown, chosen and rewarded across trials, and to reduce the size of the state space by 18^12^ states, which also made a tabular form of the model tractable. The model of the task was in epoch time (i.e. event based), rather than true time (i.e. seconds based).

The state space can be considered as a graph with edges and nodes, where the states are defined by the possible combinations of these features, and the edges are the transitions to future states. A trajectory through the state space represents one trial and as the model proceeds through a block, the trajectory traverses through the state space.

The state value, sometimes called state utility, *u*(*s*_*t*_), was calculated using the equation:

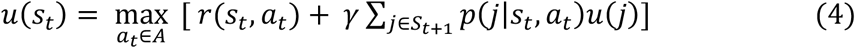

where *s*_*t*_ is the state, *a*_*t*_ is the action taken at that state, *u*(*s*_*t*_) is the state value, *r*(*s*_*t*_, *a*_*t*_) is the immediate reward, *γ* is the discount factor, *p*(*j*|*s*_*t*_, *a*_*t*_) is the transition probability to future state *j* and *S*_*t+1*_ is the set of immediate future possible states from state *s*_*t*_. A range of discount factors (*γ*=0.8, 0.85, 0.9, 0.95, 0.99, 0.999) were tested. *γ*=0.999 produced the least error for the regressions and was used for all analyses. We used value iteration to fit the MDP (Puterman, 2014). The algorithm loops over all possible states and recomputes Eq. 4 until both the policy and state values converged. This took approximately 100 iterations across the state space for each MDP that was fit.

The transitions between states include: fixation to the six cue states, cue state to token update, token update to cashout or intertrial interval and intertrial interval to fixation. The transition probabilities from fixation to any cue state were modeled as p_cues_=1/6 a there were 6 possible cue pairs. The transition probabilities from token update to cashout were p_cashout_=0 for TSCO 1-3, p_cashout_=0.33 for TSCO 4, p_cashout_=0.50 for TSCO 5, p_cashout_=1.0 for TSCO 6. Transition probabilities for the transition to a change in tokens given a cue image selection, i.e. p(change in tokens | image choice in a given condition), were fit using the behavioral data and average cue values that were extracted from the RW-RL model fits. Thus, this is not an ideal observer estimate, but rather our inference of the monkey’s estimate. These transition probabilities represent the monkey’s mapping of individual cues to outcomes, i.e. which picture leads to +2 tokens. The cue values are related to this mapping, thus making the RW-RL values an approximation to the process by which the animal learns the outcomes related to each cue image (**Fig. 2**).

We had to infer the monkey’s estimate of the number of tokens they would receive when they chose a given option. For example, in the first trial of a new block, the monkeys had no experience with any options, and therefore they should assume that choice of any option could lead to either -2, -1, 0, 1, or 2 tokens. However, after 10 trials the monkeys had a reasonable estimate of the token outcomes associated with each option. This process makes the MDP have partially observable states, as we estimate the transition probabilities using mean values from the RW-RL model. We carried out this estimate in two steps. First, we calculated the value estimates for each option, as a function of the number of trials they had seen each option using the RW-RL algorithm value estimates (Fig. 2). We then used these estimates to calculate the posterior probability that choice of a given cue would lead to a given outcome (i.e. Δ*tk*). For the outcome of no tokens, *p*(Δ*tk* = 0) = 0.25. For all other possible outcomes, the following equations were used:

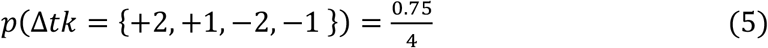

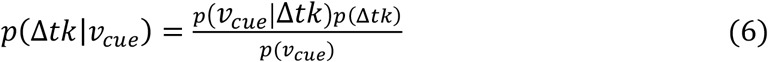

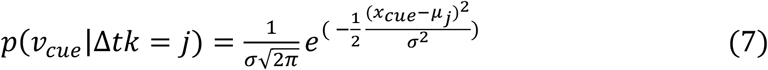

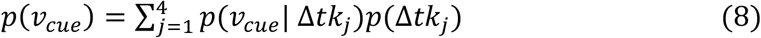

and where *v*_*cue*_ is the mean value of a single cue for a given NObs, Δ*tk*= +2, +1, -1, -2, *x* is the mean of the cue value of the chosen option, and *μ* is a mean value of one of the other cues. Transition probabilities were calculated for all possible choices and NObs and were not dependent on other MDP features such as NTk or TSCO. For example, 3 trials into the block, mean cue values were 0.17, 0.11, -0.01, -0.01 for the +2 cue, +1 cue, -1 cue, and -2 cue, respectively (**Fig. 2B**). To calculate *p*(Δ*tk* = +2|*v*_*cue*_ = 2) (i.e. the probability of receiving 2 tokens for choosing the +2 cue), *x*=0.17, *μ*_1_=0.11, *μ*_2_= - 0.01, *μ*_3_=-0.01, which produces *p*(Δ*tk* = +2|*v*_*cue*_ = 2)=0.24 in the +2 versus +1 condition. Later in the block, for example on NObs=16, mean cue values were 0.66, 0.36, -0.03, -0.04 for the +2 cue, +1 cue, -1 cue, and -2 cue, respectively. At this point in learning, *p*(Δ*tk* = +2|*v*_*cue*_ = 2)= 0.52 in the +2 versus +1 condition.

The mean cue values were used to fit the set of transition probabilities p(Δtk|choice) for Δtk= 0, 1, 2, -1, -2 and choice= cue 1, cue 2. First, an MDP was fit using the mean cue values for each animal in order to compute a converged policy of choices and action values without any free parameters. These MDP models captured general trends of behavior to select the better options (i.e. an optimal MDP) but showed faster learning than the animals learned. To better match animal learning behavior, we optimized the transition probabilities underlying the behavioral performance of the MDP.

To optimize the set of transition probabilities p(Δtk|choice), mean cue values were used with two free parameters: (1) a scaling parameter for the mean cue values such that:

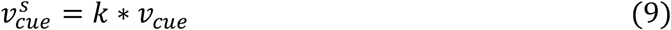

for all mean cue values and (2) an inverse temperature parameter for the choice probability (β^MDP^). In addition, the variance on the mean value curves for each animal was set to the average variance across all four cues, allowing variance to vary across trials, but not across cue values. Using the initial MDP fit and the behavioral data from the task for each animal, the two free parameters were fit jointly by minimizing the error between the MDP choice probability and average performance across sessions for each monkey (**Table 1**). The resulting parameters and transition probabilities were then used to refit the MDP until the state values and policy reconverged.

**Table 1.** MDP free parameter values for each monkey. Two parameters were optimized to minimize the error between the MDP choice probability and the monkey’s choice behavior. The mean value scaling parameter acted as a scalar on the variance of the value curves (Eq. 9). The choice probability parameter was the inverse temperature parameter used to calculate the choice probability using action values derived from the MDP.

| Monkey | Mean value scaling parameter (k) | Choice probability parameter ($\beta^{\text{MDP}}$) |
| --- | --- | --- |
| Monkey U | 0.45 | 2.77 |
| Monkey B | 0.71 | 1.87 |
| Monkey S | 1.04 | 1.15 |
| Monkey P | 1.14 | 1.34 |
| Monkey A | 0.53 | 2.24 |

In addition, MDP models were fit using a range of discount factors (*γ* = 0.8, 0.85, 0.9, 0.925, 0.95, 0.99, 0.999) for each animal’s dataset. To determine the discount factor that produced the best fit to behavior, each *γ* was used to regress on fixation reaction times, choice reaction times, p(Abort) and to produce choice probability curves for the six conditions (see below for details on the regressions). For all monkeys, *γ* = 0.999 produced the best fits to these behavioral data metrics, and thus, *γ* = 0.999 was used for all models.

### Regressions and Statistical Analyses

Comparison of performance of the MDP and RW models at predicting choice behavior was conducted using a comparison of correlation coefficients. Correlation coefficients (r_1_, r_2_) were calculated between the average choice behavior and each model separately. The values were then Fisher-z transformed to compute a p value for a two-sided test for differences between the correlation coefficients.

State values were extracted for all trials and epochs using the MDP fits for each animal. This produced a table of states such that the value of each state was: *u*(*s*_*t*_) = *f*(*NTk, TSCO, TE, NObs*). These state utilities were used to characterize trial-by-trial relationships to reaction time to acquire fixation, choice reaction time, and trial aborts.

Mean reaction times (RT) were computed by averaging reaction times across blocks of trials and then averaging across sessions for each monkey. Scatter plots of RTs from individual sessions do not include outlier reaction times. Outlier RTs were removed using Tukey’s method: RT > q0.75+ 1.5*IQR and RT< q0.25 – 1.5*IQR, where IQR is the interquartile range.

To assess the relationship between state value and reaction times to acquire fixation, linear regression on state value was performed such that:

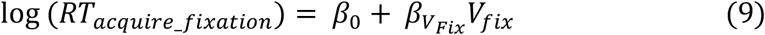

where *V*_*fix*_ is the value of the fixation state. Reaction times were log transformed before the regression.

To assess the relationship between state value and reaction times to choose, linear regression on state value was performed such that:

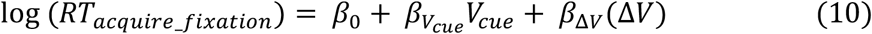

where *V*_*cue*_ is the value of the cue presentation state and Δ*V* = *V*_*cue*_ − *V*_*fix*_ . Reaction times were log transformed before the regression.

To assess the relationship between state value and the probability of aborting a trial, logistic regression on state value was performed such that:

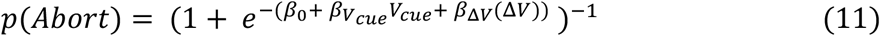

where *V*_*cue*_ is the value of the cue presentation state and Δ*V* = *V*_*cue*_ − *V*_*fix*_ . We additionally assessed the effect of cue condition on p(Abort) using a mixed effects ANOVA, where monkey was the random effect and cue condition was the fixed effect.

To assess whether regressors were significantly different than zero, for each animal, t-tests on the distributions of beta values across sessions for each regressor were performed for each animal. These values are reported in the text. Mean parameter values did not appear to be Gaussian distributed across monkeys. Therefore, to assess whether the regressors were significantly less than zero at the group level, the non-parametric Wilcoxon signed-rank test was used on the distribution of mean parameter values across animals. Results of these tests are reported in the figure captions and text. To show group trends in relationships between reaction times and regressors, 1D kernel smoothing was conducted on each monkey’s data with *σ* = 0.5 with a Gaussian kernel.

We also sought to examine the contributions of the different variables that defined the state to each regression by marginalizing over one factor at a time, to remove its effect, and carrying out the correlation analyses. To perform this marginalization, we averaged over the state values for all possible values of a single feature, given the other features. For example, to compute state values for all features averaging over NTk, with *TSCO* = 4, *TE* = 1, *NObs* = 12, we computed *V*_*fix*_= mean(*f*(*NTk* = 0: 8, *TSCO* = 4, *TE* = 1, *NObs* = 12)), as NTk=8 is the maximum number of tokens possible when TSCO=4.

## Code Accessibility

All code used to generate the results in this manuscript can be accessed on GitHub here: https://github.com/dcb4p/mdp_tokens.

## Results

Five monkeys were trained on a stochastic tokens task (Taswell et al., 2018) (**Fig. 1**). Briefly, each block of the task used four novel images, and choice of each image led to a different possible token outcome (+2, +1, -1, -2). In each trial, two of the four images were presented as options, and the monkey made a saccade to one of the cues and held their gaze to indicate their choice. After the choice, the monkey received, stochastically, the corresponding change in tokens on the screen. In 75% of the trials they received the number of tokens associated with the cue and in 25% of the trials the number of tokens did not change. Every four to six trials was a cashout trial. In cashout trials, the monkey received one drop of juice for each accumulated token. The monkey would then start over accumulating tokens until the next cashout trial. Each of the pairs of cue images (six total) were presented 18 times during a block of trials. At the end of the block, the four cue images were replaced with novel images, and the monkey restarted learning the associations between the images and token outcomes. Behavioral performance and learning were assessed for each animal by the increasing frequency with which the monkeys chose the image associated with the better option in each condition over the course of a block.

In this task the monkeys were not given rewards on every trial. Rewards were only given at the time of cashout when tokens were exchanged for juice. Commonly used RL models, such as the Rescorla Wagner model (RW) (Sutton and Barto, 1998), do not make a distinction between symbolic reinforcers and primary rewards and therefore they do not have a natural way to model the difference between primary and secondary reinforcers. To address this, we developed a state-based, Markov decision-process model of the stochastic tokens RL task to capture the relevant features of the task that would affect motivation and choice behavior. Within MDP models, values and available actions are defined by the current state. In our model, the state is a function of the number of tokens (NTk), trials since cashout (TSCO), task epoch (TE) and number of observations of each condition within the block (NObs). The state space consists of all possible combinations of these four features. The model, once trained, has states that inherit value from their proximity to the true rewarding state (cashout), similar to how a well-trained monkey expects to earn juice in the future.

State values are given by the maximum action value in each state. State values and action values are equivalent in all epochs except the choice epoch, because only one action is possible in the other epochs. Rewards are only delivered in the cashout period, and therefore immediate expected values, r(s_t_, a_t_) are 0, except in the cashout period when more than 0 tokens have been accumulated. In all other states, immediate expected values are 0, and state values are future discounted expected values, all of which are filtered through the graph from the cashout period. Thus, future expected values in each state follow from the features of the task that predict the delay to and size of the reward that will be delivered in cashout.

For example, the value of the fixation state is a combination of the number of tokens accumulated, trials since cashout, and number of observations of the cue pairs witnessed up until that trial (**Fig 3**). The value of the fixation state is the sum of the immediate expected value (which is 0 at fixation because no juice is ever delivered) and the future expected value, which is the expected value of the next state (i.e. the average over the cue states, each of which occurs with a probability of 1/6). Thus, state value in the fixation state is inherited from the values of the cue states.

**Figure 3.**
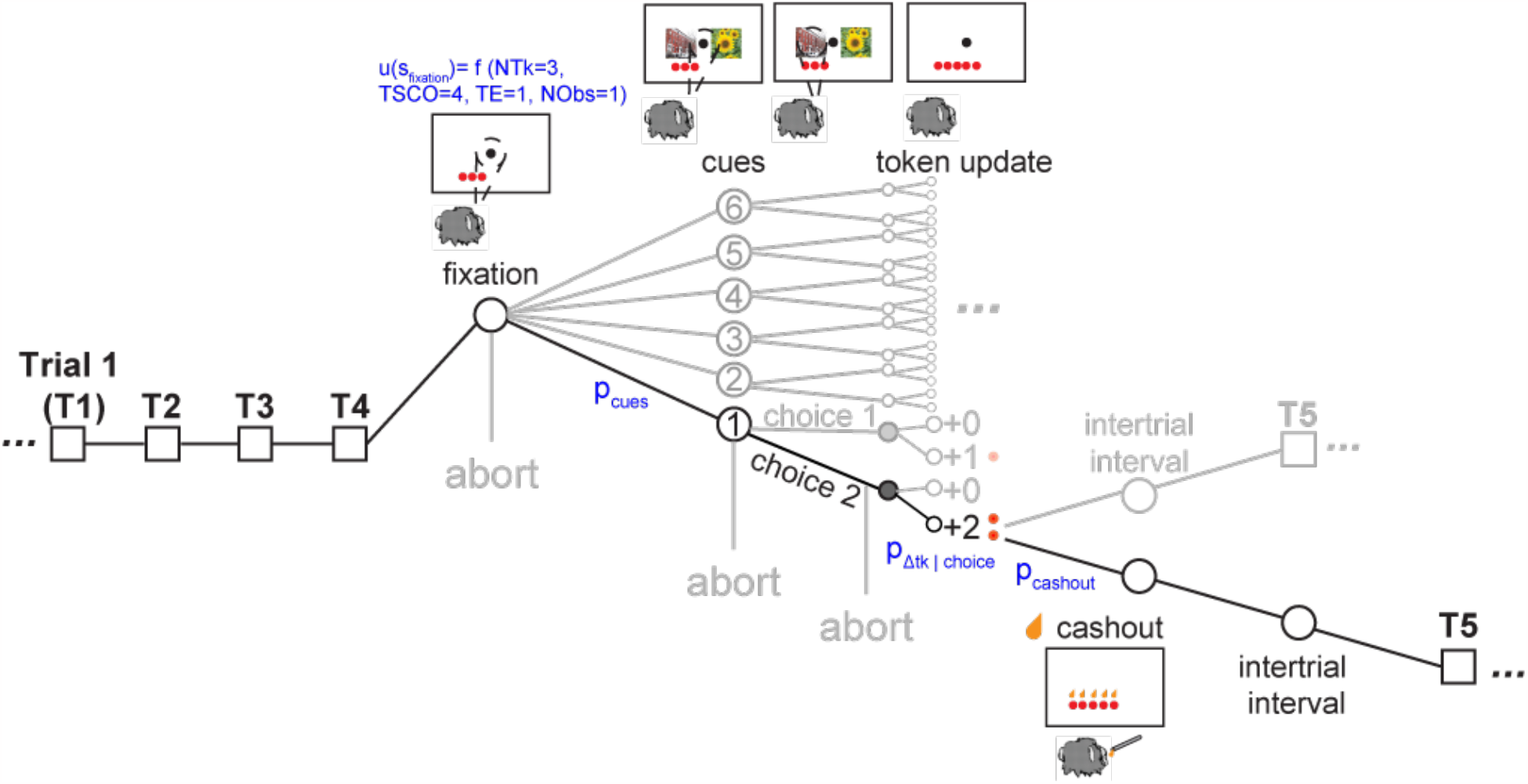
Overview of task state space for Markov Decision Process (MDP) framework. Each circular node represents a state given by a set of task features: token count (NTk), trials since cashout (TSCO), task epoch (TE), number of cue condition observations (NObs). Each edge of the graph represents a transition probability, or the transition between states that could be deterministic (p=1) or probabilistic (p <1). A single trial is highlighted and the progression from each step in the task is shown for an example trial that ends in a cashout. The example trial starts at fixation and the monkey has three tokens, making the value of the state f(NTk=3, TSCO=4, TE=1, NObs=1). The transition probability p_cues_=1/6 and represents the probability of any of the six cue conditions being the next state. At the cue states, the monkey must make a choice, and the model captures this choice policy behavior. The next set of states after the cue state is the outcome state; the transition is governed by the transition probability p_Δtk|choice 2_. In the example, the monkey receives 2 tokens and the model receives 2 tokens. After the outcome state, the monkey can transition to the intertrial interval or cashout; the transition to this state is governed by p_cashout_. During cashout, one drop of juice is exchanged for each token and all the tokens disappear before proceeding to the next intertrial interval and subsequent trial. Image credit: Wikimedia Commons (scene images).

In each cue state, there is a choice between the two cues, which then leads to the token outcome states. As five of the six possible cue conditions have gain outcomes (once the monkey learns to select the better option), the state values of the cue conditions reflect this possible gain as it develops with learning. In the outcome states the monkey can receive +2, +1, 0, -1, or -2 tokens. As there is also no immediate reward available during the cue states, state value comes from the future expected value of the intertrial interval or cashout states.

The monkey learns to choose options that maximize gaining tokens over losing tokens. We modeled this learning as an inference over the token outcome distribution associated with each choice, using a parameterized function of the number of times an option had been chosen. The model, which generated an estimate of the transition probabilities from cue to token outcome (p_Δtk|choice_), was fit to each monkey’s choice behavior. Unlike the transition from the fixation state to the cue state (p_cue_), the probability of transitioning to each outcome state (p_Δtk|choice_) changes as the monkey learns the associations between the cue options and token outcomes. For example, at the start of a block, the monkey does not know which cue image predicts which token outcome, and p_Δtk=+2|choice_ = p_Δtk=+1|choice_ = p_Δtk=-2|choice_ = p_Δtk=-1|choice_. Once the monkey learns which cue image is associated with +2 tokens, they are more likely to select that option, and we assume they infer that the probability of getting two tokens, p_Δtk=+2|choice_, is larger given choice of that option. Thus, the transition probabilities change over the course of a block as the number of times the cue pair has been observed (NObs) increases. The changes in these transition probabilities reflect learning (Fig. 2B,C).

At the time of token outcome, the next possible states are either the intertrial interval or cashout, governed by the transition probability p_cashout_. This transition is a feature of the task and does not change with learning. At the time of cashout, the state value is the sum of the immediate reward (1 drop of juice per token present) and the next state, which is the intertrial interval, with zero tokens. In the model, this means state value will drop after cashout if the monkey cashed out tokens for juice, as the state value depends on the number of tokens.

### Changes in task features drive fluctuations in state value

State values change with variation of each state feature. For example, having more tokens increases state value (**Fig. 4A**). Being closer to the cashout state (e.g. TSCO >4) also increases state value (**Fig 4B**). As the monkey proceeds through the task epochs (i.e. fixation, cue onset, outcome, cashout or intertrial interval), the state value will also increase for conditions where tokens can be gained, and more subtly, because one is also getting closer to cashout (**Fig. 4C**). For the -2 v. -1 cue condition where tokens can only be lost or maintained (if there is a no change outcome), the value of the state either decreases or increases marginally approaching cashout (**Fig 4D**). In the first trial of a block, when the monkey has started learning the associations between cues and outcomes (i.e. NObs =1), the best option in a pair of cues will be ambiguous. This is reflected by identical state-action values in the model (**Fig. 4E**). Near the end of the block, when the monkey knows which cues correspond to +2 and +1 tokens (e.g. NObs 18), the state action values will reflect the knowledge of the best option and the state values at the time of the cue state will be higher for the conditions with +2 or +1 cues (**Fig. 4E**). Even though the fixation state precedes the cue state, the number of observations also affects the value of the fixation state and causes it to increase as NObs increases, because the monkey can make better choices when the options are presented (**Fig. 4F**). The minimum state value is at the baseline for all features, i.e. NTk=0, TSCO=1, TE=Fixation, NObs=1 (**Fig. 4F**, NObs=1).

**Figure 4.**
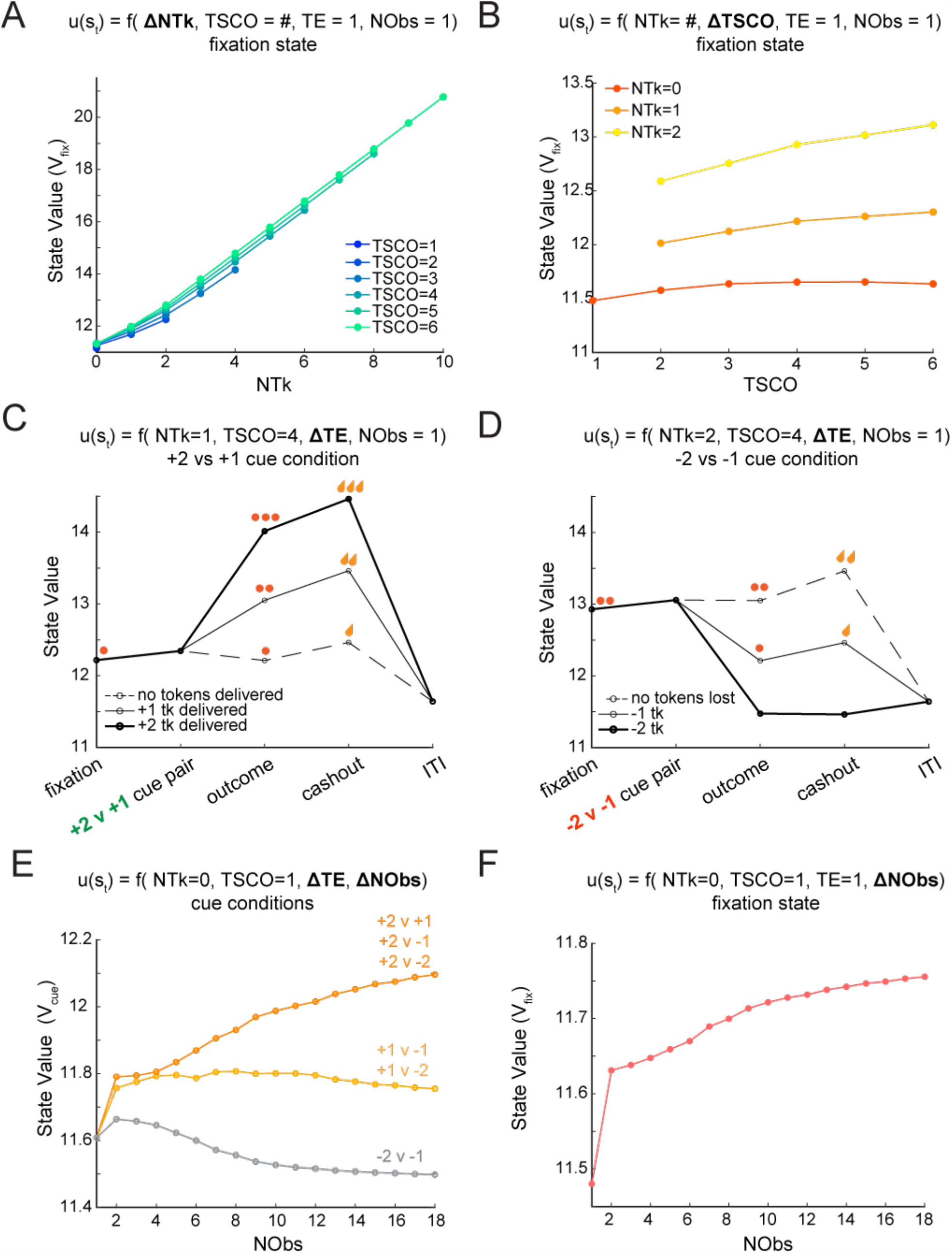
How changes in state features can affect state value (example from Monkey B). **(A)** Fixation state value (V_fix_) versus token count (NTk). As NTk increases, state value increases. For one trial since cashout (TSCO=1), only 1 trial has been completed, so the maximum number of tokens available is two. As the TSCO increases, juice will be delivered sooner, which is reflected by higher state value. **(B)** Fixation state value (V_fix_) versus trials since cashout (TSCO). As TSCO increases, the state value increases when there is more than one token (NTk=1, NTk=2). If there are zero tokens (NTk=0), then state value increases until TSCO=4. When cashout becomes possible, state value begins to decrease when there are no tokens, because a cashout epoch can occur, with zero juice delivery, resetting the interval before juice can be delivered again. **(C)** State value versus task epoch for a single cue condition +2 vs +1. As the model proceeds through a trial, state value changes depending on task epoch and outcome. In this example, there is one token at fixation (red dot, far left) and three traces are shown, one to represent each possible outcome (+2 tokens, +1 token, 0 token change). State value increases for gaining tokens and decreases slightly when no tokens are gained. State value at cashout corresponds to the token count. The state value is identical for all three outcome traces during the intertrial interval after a cashout. **(D)** State value versus task epoch for a single cue condition -2 vs -1. In this example, there are two tokens at fixation (red dots, far left) and three traces are shown, one to represent each possible outcome (-2 tokens, -1 token, 0 token change). State value decreases when tokens are lost and stays constant when the outcome is zero tokens. At the time of cashout, state value depends on whether tokens are present. Like in (C), the state value is identical for all three outcome traces during the intertrial interval after a cashout. **(E)** Cue state value (V_cue_) versus number of observations of a cue pair (NObs). As NObs increases, the state value increases for all cue conditions that include the best option (+2 tokens). The state value for the cue conditions with the +1 option decreases with learning and plateaus. The state value for the loss vs. loss condition (-2 v -1) decreases with NObs. **(F)** Fixation state value versus NObs. As NObs increases, the value of the fixation state increases. As the monkey proceeds through a block, they learn the associations between the cue images and token outcomes, and it is more likely the monkey will select the better options (+2 and +1). In the MDP, this means that as NObs increases, it will be more likely that tokens will be received, which causes an increase in the future expected value and thus state value.

The exact value of the baseline state value and the relationship between NObs and the fixation state value vary by model (i.e. monkey). This relationship is affected by three things: the token outcome transition probabilities, the discount factor, and number of iterations for fitting the model. The discount factor was 0.999 for all monkeys, and the number of iterations for fitting each model was constant. Only the token outcome transition probabilities (p_Δtk|choice_), which were fit to each monkey’s behavior, vary between monkeys in the models. Therefore, the larger the token outcome transition probabilities to gain outcomes, the larger the initial state value even at the time of fixation. In other words, when the monkey learned faster, these transition probabilities changed faster, and state value increased faster with NObs.

### State-based MDP model of symbolic reinforcement captures learning behavior

To test the validity of the choice policy of the MDP model for each monkey, we calculated the choice probabilities produced by the choice policy of each MDP, after passing action values through a softmax. After fitting an MDP to each monkey, choice probability was calculated using the action values for each choice in each cue condition, for each trial in a block (NObs 1-18). The action values were passed through a softmax with an inverse temperature parameter β (see methods), which controlled the stochasticity of the choice policy given two action values. Average choice probability across animals demonstrated that the choice probabilities produced by the MDP produced similar fits to the behavioral data to the RW-RL model, with no statistically significant difference between the correlation coefficients computed from the behavioral data and the two models (r_RW_= 0.9904, r_MDP_= 0.9868, difference in correlation: p= 0.25, **Fig. 5**). This verified that the MDP captures choice behavior in the task, over and above its ability to model future state values. Further analyses using state values are, therefore, grounded in an accurate representation of choice behavior.

**Figure 5.**
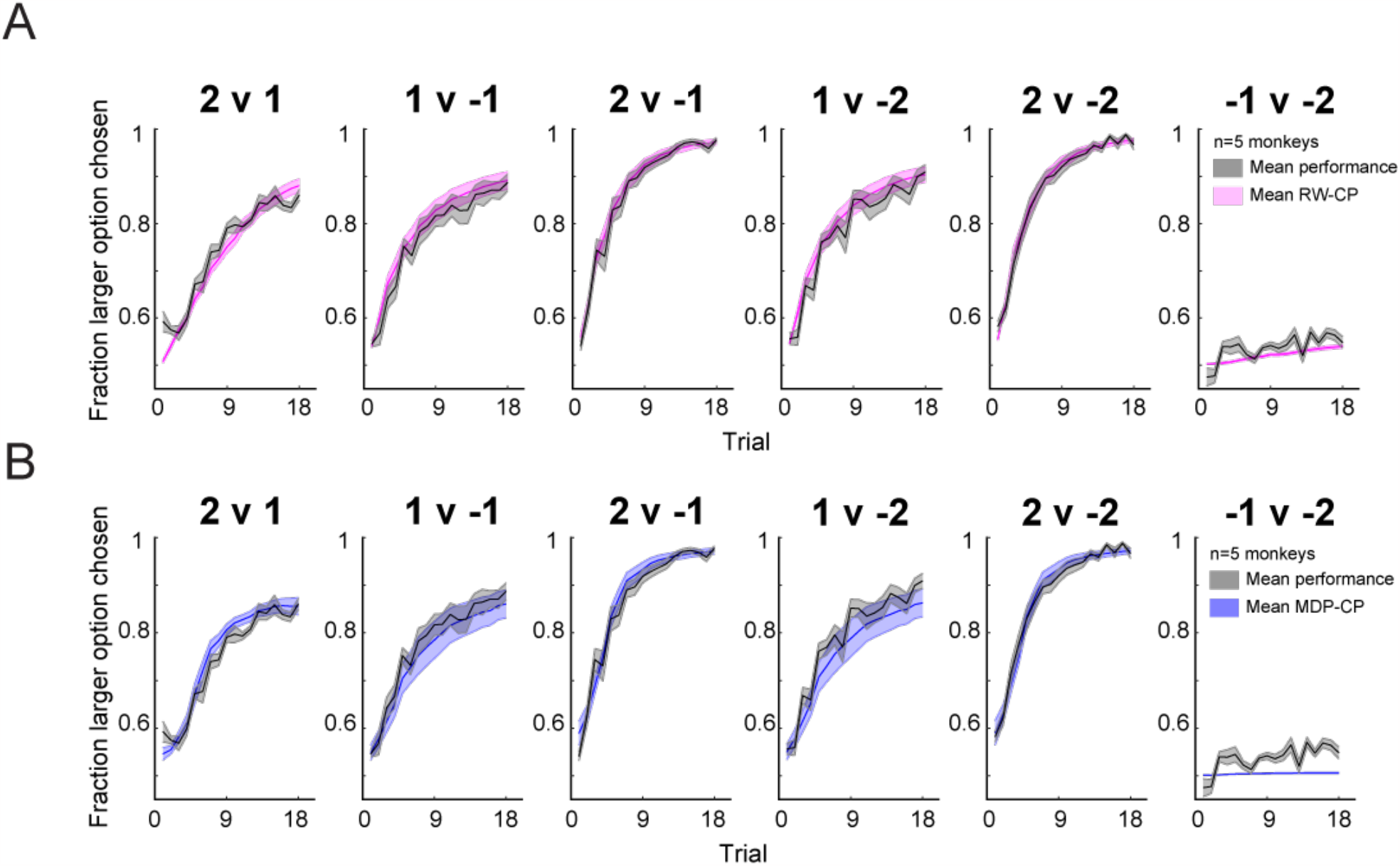
Average performance in the tokens task and model fits. **(A)** Average performance and RW-RL model fits for all subjects (n=5) monkeys in the six task conditions (s.e.m. across animals). **(B)** Same as (A) but average MDP choice probability instead of RW-RL model fits.

### Time to acquire fixation is related to the value of the fixation state

We next examined whether the MDP state values could be used to predict motivation in monkey behavior. The first question was how state value might affect the initiation of a trial, which has been previously shown to be affected by motivation (Hamid et al., 2016; Oemisch, Johnston and Paré, 2016; Mohebi et al., 2019; Steinmetz et al., 2019). For example, if the monkey has multiple tokens at the start of a trial, might they be more motivated to initiate a trial, than in the case when they have no tokens (**Fig. 6A**)? However, token count is not the only task feature that could affect motivation in this task. Thus, we used the value of the fixation state (V_fix_) from the MDP to relate all relevant task features (NTk, TSCO, TE, NObs) on a trial-by-trial basis to the time it took the animal to acquire the fixation spot. We conducted a linear regression for each session of data from each animal and calculated the average regression coefficient value for each animal (**Fig. 6B**). Mean regression coefficients for V_fix_ (β_Vfix_) were significantly less than zero at the group level (p= 0.0312, Wilcoxon signed-rank test). All five of the individual distributions of regression coefficients for each animal were statistically significant (t(20)= -5.78; t(22)= -9.48; t(16)= -11.74; t(19)= -28.18; t(19)= - 19.37; p<0.0001). Thus, as the value of the fixation state increased, reaction times decreased (**Fig. 6C**) and this was true for all five monkeys (**Fig. 6B, 6D**).

**Figure 6.**
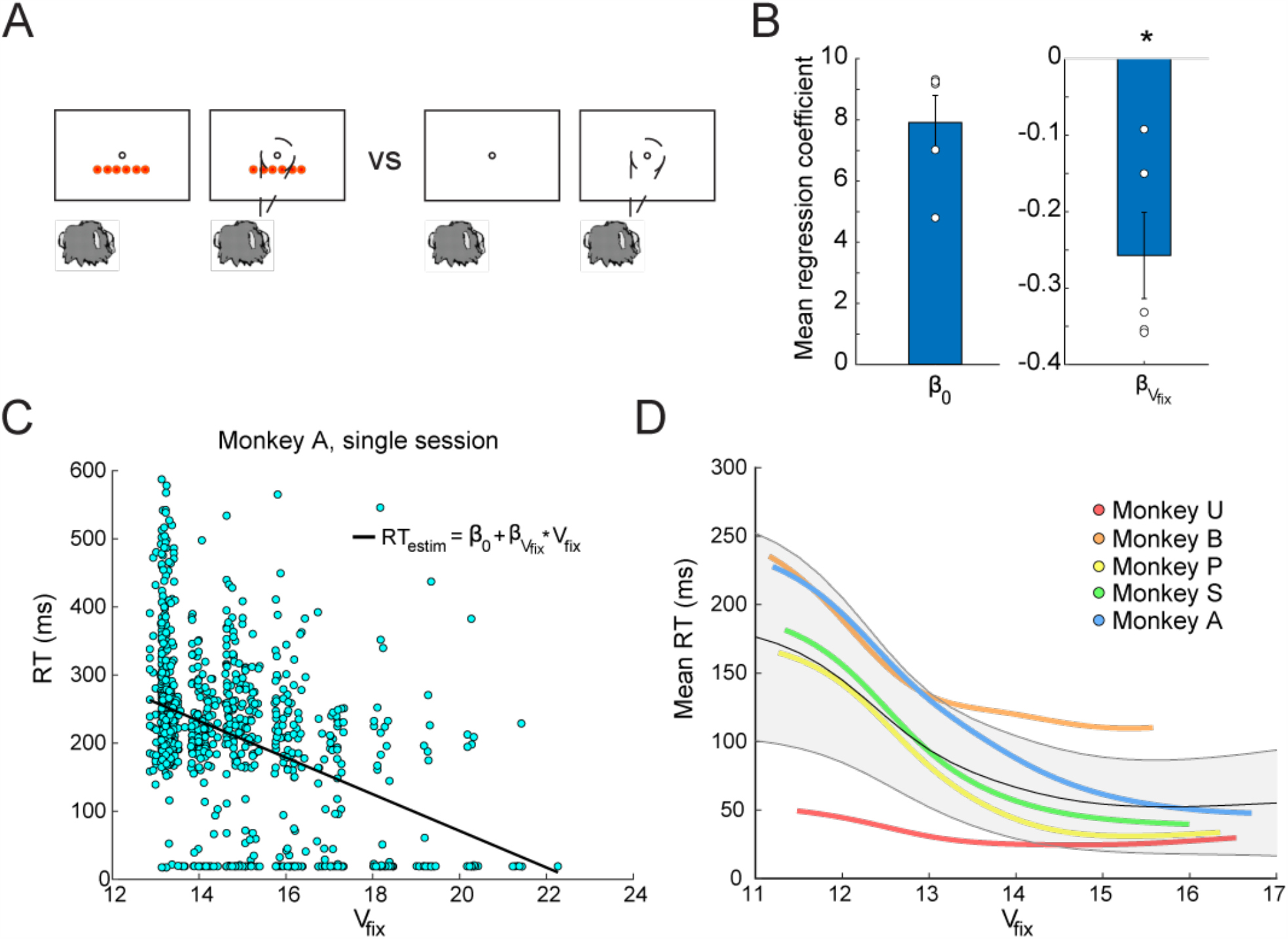
Reaction time to acquire fixation. **(A)** An example case of when motivation might differ in the tokens task at the time of fixation. **(B)** Mean regression coefficients across animals from the linear regression on V_fix_ (bar plot) and mean beta values across sessions for each animal (dots). (* indicates p< 0.05). Note that regressions were conducted on log(RT). **(C)** An example set of reaction times from a single session from Monkey A showing a decrease in reaction time to acquire fixation as V_fix_ increases with an overlay of the regression line. Values near zero indicate trials in which the monkey was already within the fixation window when the fixation cue appeared. **(D)** Kernel smoothed, averaged mean reaction times for each monkey versus V_fix_. Average reaction times across sessions are shown for each animal in a different color indicated by the legend. The average of all animals is shown in grey, with error bars showing the standard deviation across animals.

### Choice reaction time is related to the value of the cue state and change in state value

Next, we asked whether choice reaction times were related to the value of the cue state (V_cue_) and the change in value between the cue onset and fixation state (ΔV= V_cue_ - V_fix_), which would reflect an impending gain or loss of tokens from selecting a cue. For example, if the cue condition was loss v. loss (-1 vs. -2), the monkey might be slower to choose an option than in the case of gain v. gain (+1 vs. +2) where there is a preferred option (**Fig. 7A**). We conducted a linear regression of V_cue_ and ΔV on reaction times for each session of data from each animal and calculated the average regression coefficient for each animal (**Fig. 7B**). Mean regression coefficients β_Vcue_ and β_ΔV_ were significantly less than zero at the group level (p= 0.0312, Wilcoxon signed-rank test). This meant that as the value of the cue state increased, reaction times became faster, and as the change in cue value became more positive, reaction times also became faster (**Fig 7C, 7D**). Four of the five individual monkey distributions of β_Vcue_, where session was the repeat, were statistically significant (t(20)= -2.19, p <0.05; t(22)= -6.62, p <0.0001; t(16)= -3.44, p <0.005 ; t(19)= -1.98, p =0.06 ; t(19)= -14.78, p <0.0001). Five of the five individual distributions of β_ΔV_ were statistically significant t(20)= -7.29, p <0.0001; t(22)= -8.86, p <0.0001; t(16)= -3.22, p <0.01 ; t(19)= -3.89, p <0.001 ; t(19)= -9.98, p <0.0001). In summary, this demonstrated that as the value of the cue state was higher, choice reaction times were faster for all animals. These analyses also demonstrated that when the change in state value from fixation to cue (ΔV) was positive, reaction times were also faster.

**Figure 7.**
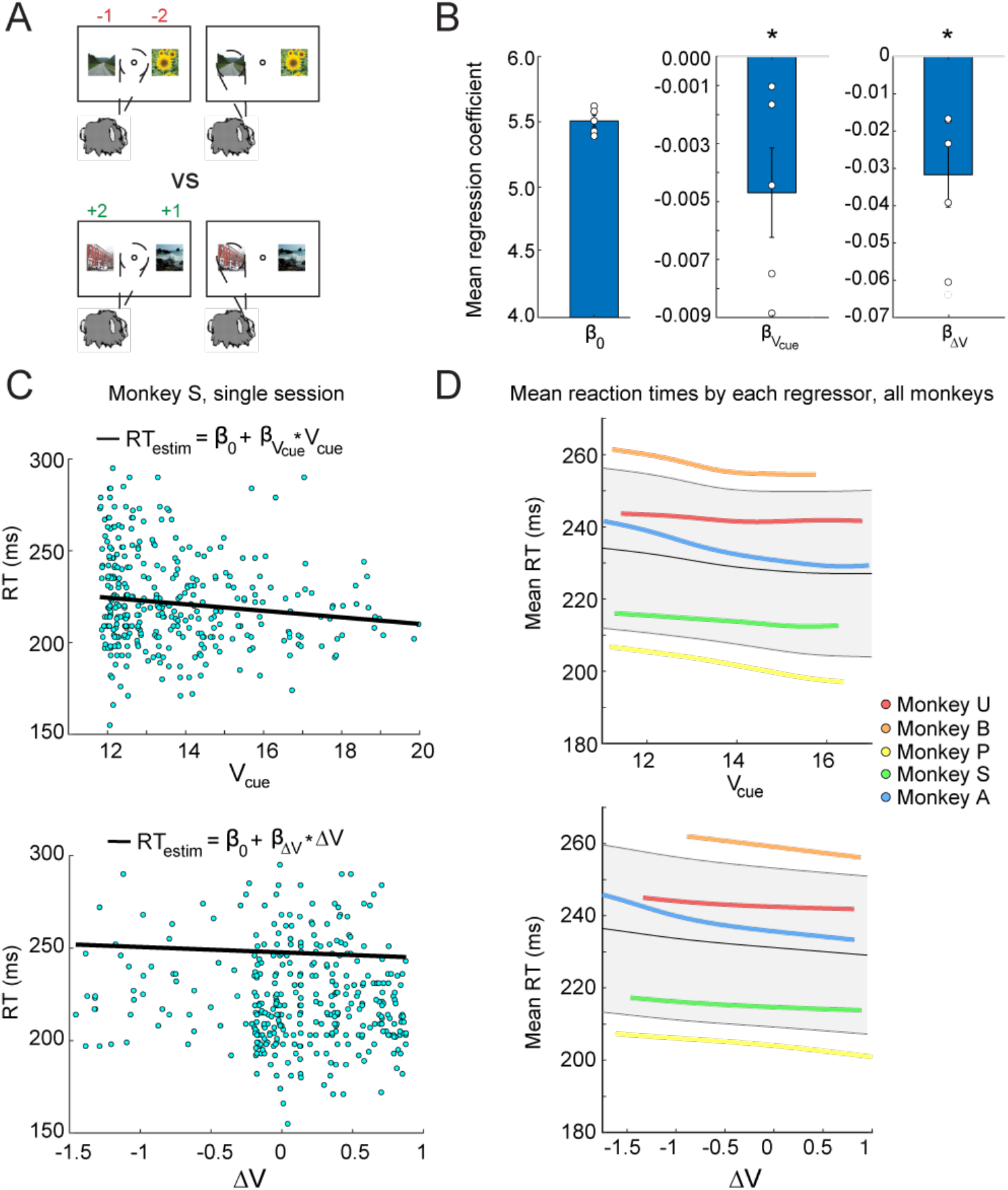
Reaction time to choice. **(A)** An example case of when motivation might differ in the tokens task at the time of image cue presentation. Image credit: Wikimedia Commons (scene images). **(B)** Mean regression coefficients cross animals from the linear regression on V_cue_ and ΔV (bar plots) and mean beta values across sessions for each animal (dots). (* indicates p< 0.05). Note that regressions were conducted on log(RT). **(C)** Example set of reaction times from a single session from Monkey S showing a decrease in choice reaction time as V_cue_ increases and ΔV becomes more positive with regression fits overlayed. **(D)** Kernel smoothed, averaged mean reaction times for each monkey versus V_cue_ and ΔV. Average reaction times across sessions are shown for each animal in a different color indicated by the legend. The average of all animals is shown in grey, with error bars showing the standard deviation across animals.

### The probability of the monkey aborting a trial is related to the value of the cue state

To investigate the relationship between the completion of a trial and state value, we related the frequency of trial aborts to state value by looking at all trials in a session and analyzing both complete and incomplete trials. If the monkey moved his eyes outside the fixation window during fixation, did not choose a cue, or did not hold the cue long enough, the trial was aborted and repeated. Given that the monkeys do not learn to pick the smaller loss well in the loss vs. loss condition (Fig. 5), it might be more likely that the animal aborts these trials to avoid losing tokens (**Fig. 8A**). Indeed, past work has shown that monkeys are more likely to abort cue conditions with two loss cues (Taswell et al., 2018). We found a significant effect of cue condition on the frequency of aborts (mixed effects ANOVA, main effect: cue condition F(5, 29)=7.86, p<0.001; random effect: monkey F(4,29)=51.14) (**Fig. 8B**). We next asked whether cue state value and changes in state value were related to the probability of aborting a trial by conducting logistical regression on cue state value (V_cue_) and the change in value between the cue state and fixation state (ΔV= V_cue_ - V_fix_). The distribution of mean regression coefficients β_Vcue_ was significantly less than zero at the group level (p= 0.0312, Wilcoxon signed-rank test) whereas β_ΔV_ did not emerge as significant (p=0.4062, Wilcoxon signed-rank test), suggesting that changes in state value were not the main factor related to abort behavior (**Fig. 8C**). Four of the five individual distributions of β_Vcue_ were statistically significant (t(20)= -3.33, p <0.01; t(22)= -19.78, p <0.0001; t(16)= -3.46, p<0.01; t(19)= - 1.94, p =0.068 ; t(19)= -14.78, p <0.0001). Overall trends across animals showed that as the value of the cue state increased, the probability of aborting decreased (**Fig. 8D**).

**Figure 8.**
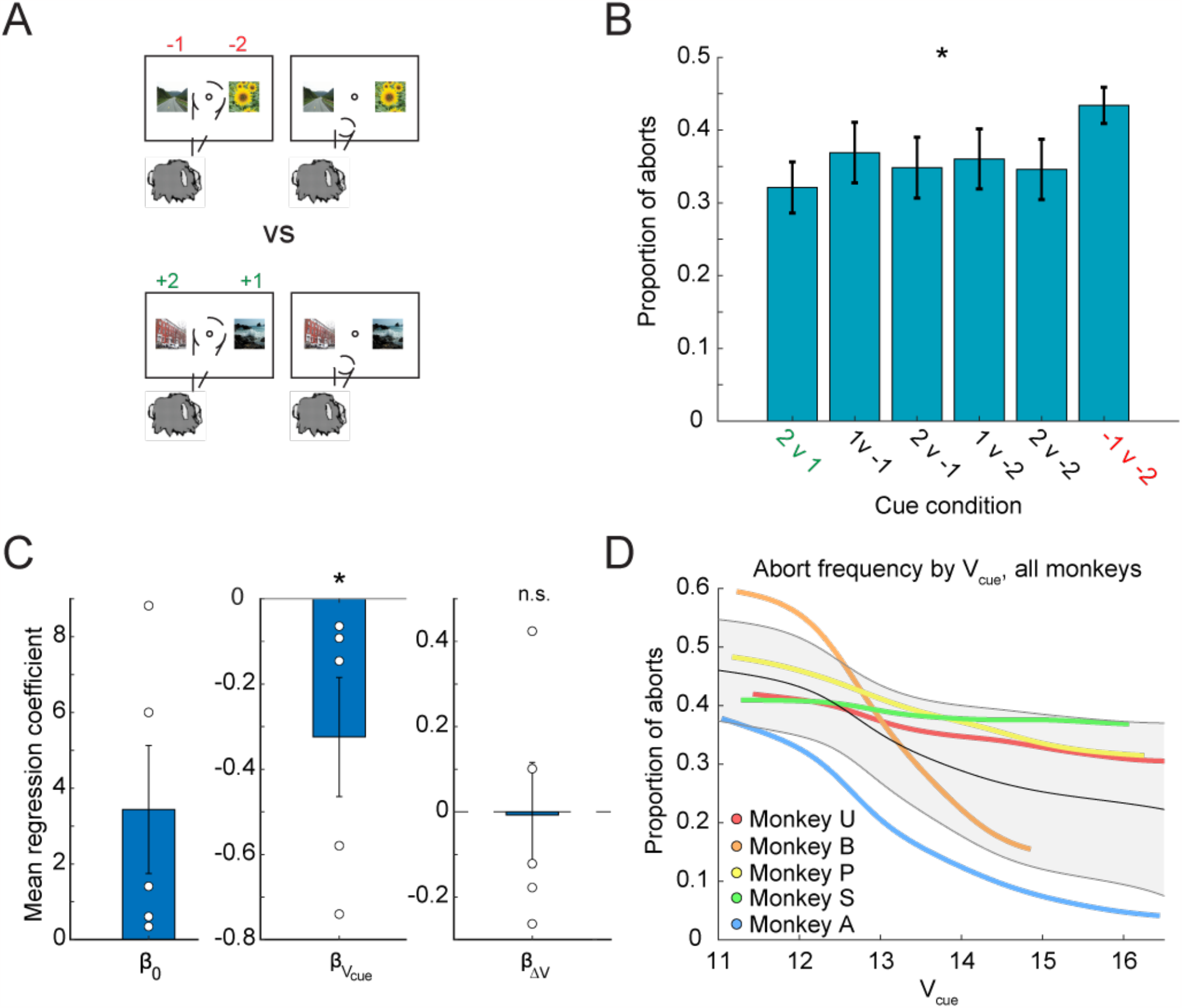
Probability of aborting a trial. **(A)** An example case of when motivation to complete a trial might differ in the tokens task at the time of image cue presentation. Image credit: Wikimedia Commons (scene images). **(B)** Average proportion of aborts in each task condition (s.e.m. average across n=5 monkeys, * indicates p< 0.05). **(C)** Mean regression coefficients across animals for the logistic regression on V_cue_ and ΔV. Only the regressor for V_cue_ was statistically significant. **(D)** Kernel smoothed, averaged mean reaction times for each monkey versus V_cue_. Average proportions of aborts across sessions are shown for each animal in a different color indicated by the legend. The average of all animals is shown in grey, with error bars showing the standard deviation across animals.

### All MDP state features contribute to state values

We next examined whether token count was the only driving force for the correlations found between state values and behavior. As was shown, current token count strongly influenced state value (Fig. 4). To assess the contribution of each of the features in the MDP to the regression results, we marginalized across each feature, thus removing the effect of variation in that feature on state value, and recomputed the regressions. For example, to marginalize over token count, state values for each trial were extracted using only the other features (TSCO, TE, NObs) after averaging over the values for all possible values of token count. The average of these state values was used as the single trial state value for the regressions.

For reaction time to acquire fixation, all distributions of regression coefficients remained statistically significant for each marginalized version of the regression (**Fig. 9A**). Removing token count from the regression had the largest effect on reducing the relationship between V_fix_ and reaction time to acquire fixation. Removing TSCO and NObs in the regression for reaction time to acquire led to an increase in beta values, which suggests these factors interact and affect the regression, but are less important than token count in the regression. For both choice reaction time and probability of aborting a trial, regressions were recomputed using only one regressor, for V_cue_. This was because removing the cue condition from the regression caused ΔV=V_cue_-V_fix_ to go to zero and therefore made the regressions uninterpretable. Marginalizing over cue condition or tokens in the regression for choice reaction times reduced the magnitude of the regression coefficients (**Fig. 9B**). This reflects a weaker relationship between the state value and reaction times without these features. In the logistic regression for aborts, marginalizing over tokens also had the largest effect on the regressors, but did not eliminate the relationship between state value and the probability of aborting a trial (**Fig. 9C**). Taken together, these analyses show that the number of tokens strongly affects all behavioral measures but is not the only factor leading to the relationships between behavior and state value.

**Figure 9.**
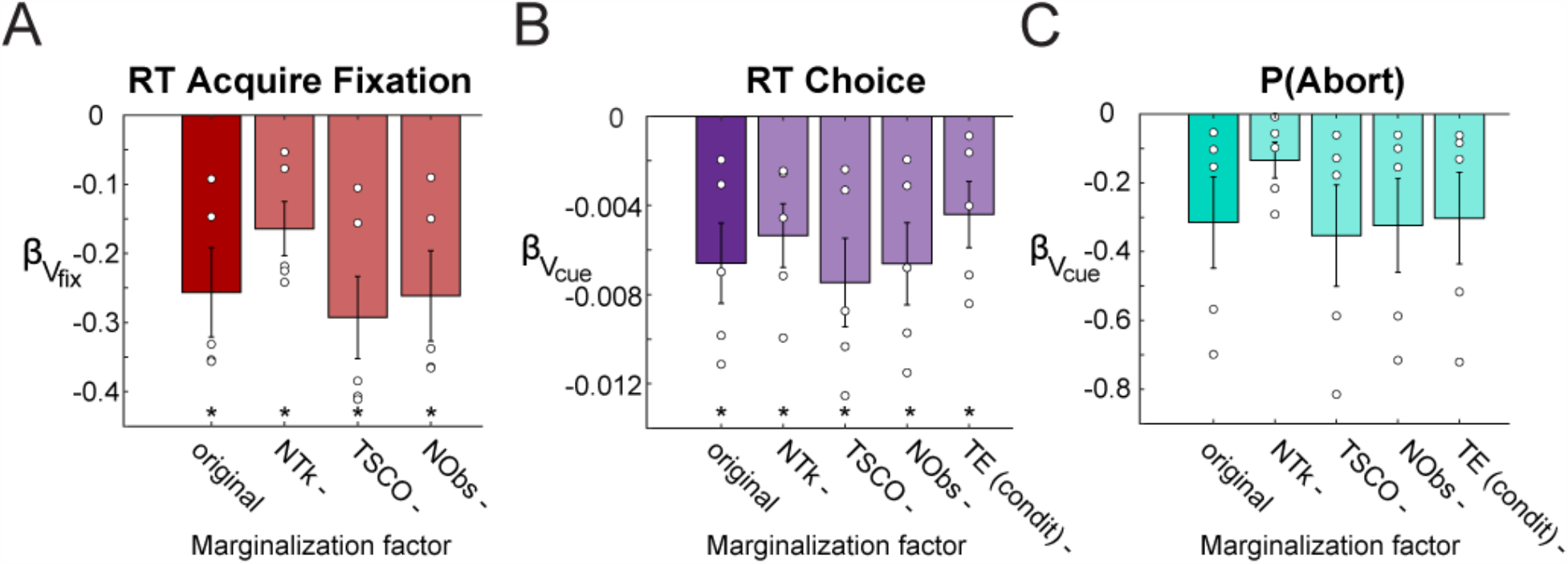
Marginalization over features. Linear regressions for each behavioral feature were recomputed using state values that omitted the effect of a single feature at a time: number of tokens (NTk), trials since cashout (TSCO), number of observations of a cue pair (NObs), Task Epoch (TE, condit). Mean regression coefficients across animals are shown (bar plots) and for each subject (dots) for three behavioral features: **(A)** RT to acquire fixation **(B)** RT to choice **(C)** Probability of aborting trials. Error bars s.e.m.

## Discussion

In this study, monkeys learned to make choices to maximize gains and minimize losses of tokens. The tokens were symbolic reinforcers that represented future juice rewards. We designed a Markov Decision Process (MDP) model to capture the relationship between features of the task that drive behavior (i.e. states) and value. We then related these state values to measures of motivation. The state space for the task included the number of tokens (NTk), trials since cashout (TSCO), task epoch (TE), and the number of observations of each cue pair (NObs). We found that reaction times to acquire fixation, choice reaction times, and the probability of aborting a trial were significantly related to state value and changes in state value (except abort probability). Furthermore, we demonstrated that state values were dependent on all state features, not just the number of tokens. Number of tokens did, however, often have a large effect. These relationships between state value and behavior cannot be captured by simpler models such as the Rescorla-Wagner model, as these models are stateless and therefore cannot capture state values that depend on future rewards, nor can they account separately for tokens vs. primary rewards. Given that the MDP also allows for modeling trial state-dependent values, it can also be used in future work to understand the neural circuitry relevant to the task.

Past work has shown that symbolic (or secondary) reinforcers can drive learning and have motivational properties similar to those of primary rewards (Wolfe, 1936; Wyckoff, 1959; Jackson, 1996; Sousa and Matsuzawa, 2001; Seo and Lee, 2009; Donahue and Lee, 2015; Farashahi et al., 2018; Taswell et al., 2018; Beran and Parrish, 2021; Yang, Li and Stuphorn, 2022; Taswell et al., 2023). This happens through the learned associations between tokens and primary reinforcers. In this task, tokens and cue images both predict rewards, although in different ways. Cues are stochastically linked to tokens on short timescales, whereas tokens are deterministically linked to juice on longer timescales. Cues, therefore, predict rewards, but only through tokens. The cues also change in each block, which requires rapid learning of the cue values, whereas the relationship between tokens and juice is stable and constant over the course of the experiment. The state-based modeling framework presented here accounts for the differential attributes of cues and tokens and allows for examining behavioral measures related to motivation, including trial initiation time, choice reaction times, and trial aborts.

The time to initiate a trial has been studied previously as a measure of motivation (Hamid et al., 2016; Oemisch, Johnston and Paré, 2016; Mohebi et al., 2019; Steinmetz et al., 2019). In a task which required rodents to nose poke after a light went on, rodents were faster, interpreted as increased motivation, when reward rate was higher (Hamid et al., 2016; Mohebi et al., 2019). When we investigated the relationship between state value and reaction times to acquire fixation, we found that a higher state value correlated with faster reaction times to acquire fixation. This implies a somewhat counterintuitive result: that on the trials immediately after receiving reward (during cashout), when state value is lowest, the monkeys are, on average, slower to initiate the next trial. Thus, symbolic reinforcers have assumed the motivational properties of rewards to encourage the choice to begin work.

Past work on choice reaction times has also suggested that reward expectation can influence execution of a choice response (Hollerman, Tremblay and Schultz, 1998; Wrase et al., 2007). Our regressions on cue state value and changes in state value from fixation suggested that reaction times to choose options were affected by other task factors, including distance to cashout, the number of tokens present, and the desirability or value of the cue condition. This fits with past work that has shown that expected outcomes can affect reaction times (Hollerman, Tremblay and Schultz, 1998; Shidara, Aigner and Richmond, 1998). In the Tokens task, once the monkeys knew the values of the cue images, the images served as a similar instruction to the possible outcomes as in past studies. Regressions on cue state value showed that as cue state value increased, reaction times decreased, as the monkeys learned to anticipate gains from certain cue conditions. Correspondingly, in loss vs. loss (-2 v -1) trials, the monkeys slowed their choices.

Aborted trials can happen for many reasons. In our Tokens task, however, we observed a systematic increase in abort trials in the condition involving only loss options (-2 v -1), which led us to investigate how cue state value and changes in state value might correlate with this behavior. Past work on trial abort behavior has shown that aborts (or refusals) occur most often in trials furthest from reward (La Camera and Richmond, 2008; Inaba et al., 2013) and trials that require the most effort (Pasquereau and Turner, 2013, 2015; Varazzani et al., 2015), suggesting that animals are more motivated to complete a trial when the cost of reaching a reward is lower. Our regression results are consistent with these findings, as monkeys were less likely to abort when the cue state value was higher.

Our analysis showed that monkeys were motivated to work when they had more tokens. However, as our marginalization over dimensions of the state vector showed, state values and our regression results depend on more than the number of tokens present. In this task, higher state value, and therefore higher discounted future expected reward, led to faster trial initiation, faster reaction times, and fewer aborts. This has implications for understanding the neural responses, as the time leading up to the receipt of the reward, also known as the anticipatory phase (Knutson et al., 2001; Ernst et al., 2004; Rademacher et al., 2017), has signals that capture expectation of future reward, which occurs in the consummatory phase (Dillon et al., 2008; Kumar et al., 2014). Understanding the dynamics of anticipation, motivation, and reward in a single framework allows for linking both processes to fluctuations in neural activity in multiple brain areas.

Within the presented framework, symbolic reinforcers have been recast as dimensions that drive state value. Past work involving choice tasks and state value have suggested the existence of a ventral circuit for the representation of state value (Gläscher et al., 2010; Averbeck and Murray, 2020) and state transitions (Belova, Paton and Salzman, 2008; Chan et al., 2021; Kalmbach et al., 2022). It has been hypothesized that distinct ventral and dorsal networks define behavioral goals and orchestrate actions to achieve goals, respectively (Everitt et al., 1999; Cardinal et al., 2002; Averbeck and Costa, 2017; Averbeck and Murray, 2020). In choice tasks, the main behavioral goal is to reach high value states. Recent work has shown correlations between fluctuations in dopamine and state value (Hamid et al., 2016), and local control of dopamine in the ventral striatum, is related to motivation (Mohebi et al., 2019). However, the ventral circuit, which includes the orbital frontal cortex, ventral medial prefrontal cortex, ventral striatum, ventral pallidum, and amygdala, is innervated by dopaminergic projections in multiple sites (Haber, 2014), and thus dopamine may differentially affect processing in each of these areas to support reinforcement learning and motivation (Berke, 2018; Westbrook and Frank, 2018). Furthermore, recent lesion work has shown that lesions of the ventral striatum and amygdala show only subtle deficits on performance on the Tokens task (Taswell et al., 2018; Taswell et al., 2023) but larger deficits in reversal learning tasks (Costa et al., 2016) and tasks requiring switches between action-based and stimulus-based strategies (Rothenhoefer et al., 2017).

The question then becomes, how are connections between symbolic reinforcement, rewards, and actions represented in the brain? Symbolic reinforcers such as tokens could be tracked directly across multiple areas, as a global representation of visual object numerosity (Tudusciuc and Nieder, 2009; Ramirez-Cardenas, Moskaleva and Nieder, 2016; Viswanathan and Nieder, 2020), but numerosity does not directly have motivational value. However, symbolic reinforcers can take on a range of identities. Furthermore, other states including abstract completion of intermediate goals can serve as symbolic reinforcers (Janssen et al., 2022). Furthermore, as the capacity to measure more neural signals simultaneously has advanced, there has been growing evidence that task-related signals are represented across many areas (Dotson et al., 2018; Steinmetz et al., 2019; Fine et al., 2023). Therefore, it is unlikely that there would be a localized neural signature of an individual task feature, as most task features will be represented across many areas. Thus, we must consider how symbolic reinforcers might be mapped onto a distributed representation that allows for flexibility in the identity of the reinforcer, and design future experiments with this in mind. Here, we have selected four features to integrate: tokens, temporal distance to reward, task epoch, and cue observations to measure the state value moment by moment in the task. The model therefore generates values for each task state, including fixation, cue presentation, token outcome, and the inter-trial interval.

In summary, we developed a computational framework that quantifies the value of symbolic reinforcers and characterizes the effect of several task features on those values. Furthermore, the model captures not only choice behavior, but also behaviors related to motivation. In this task, reaction times to initiate a trial, choice reaction times, and the probability of completing a trial were correlated with state value and changes in state value. Our results suggest that symbolic reinforcers and rewards can have similar effects on behavior, which allows for predictions about how symbolic reinforcers might be represented in the brain.

## Acknowledgements

This work was supported by the Intramural Research Program of the National Institute of Mental Health (ZIA MH002928 (BA)). The funders had no role in study design, data collection and analysis, decision to publish, or preparation of the manuscript.

## Notes

### Competing Interest Statement

The authors have declared no competing interest.

